# GEF me a break: the consequences of freezing Rho guanine-nucleotide exchange factor catalytic domains

**DOI:** 10.64898/2026.04.08.717323

**Authors:** Lauren K. Anderson, Emma Barpal, Herra Mendoza, Jennifer N. Cash

**Affiliations:** Department of Molecular and Cellular Biology, University of California — Davis, Davis, CA, USA

## Abstract

Purified proteins are routinely flash frozen for use in functional and structural studies, providing a convenient way to reproduce results across complex experiments. Rho guanine-nucleotide exchange factors (RhoGEFs) are no exception to this practice, yet the effects of freezing on their activity and stability remain largely uncharacterized. This gap potentially affects the characterization of these important enzymes and how results are interpreted with respect to their prospective use as therapeutic targets. Here, we tested the isolated DH/PH tandems of P-Rex1, P-Rex2, and PRG under different cryoprotectant conditions and monitored activity and thermostability over time after flash freezing. Our results show a clear divergence between the activity of fresh and frozen purified RhoGEF protein samples in as little as one week for some conditions. Specifically, the variability in data collected on frozen samples was greatly increased. Despite these differences, thermostability seems to be preserved for much longer timepoints across RhoGEFs. Moreover, despite eventual changes in both activity and thermostability with respect to freezing, there are no obvious changes in global conformation between fresh and frozen samples of the isolated P-Rex2 DH/PH tandem. From our data, there are few generalizable trends between the different RhoGEFs and no single cryoprotective agent tested was a silver bullet to preserve both activity and thermostability across RhoGEFs. Overall, our findings emphasize the unpredictable effects of freezing RhoGEFs. As such, RhoGEF freezing should be carefully characterized for each protein and critically viewed when comparing analyses between different studies.

## Introduction

Dbl Rho guanine-nucleotide exchange factors (RhoGEFs) are important regulators of small Rho GTPases which control cell movement and proliferation (Narumiya 1996; Bourne et al. 1990). Common across most Dbl RhoGEFs is a tandem catalytic Dbl homology (DH) domain and pleckstrin homology (PH) domain which are structurally well-characterized (Fig. S1). The DH domain fold is composed of six α-helices forming an oblong helical bundle (Rossman et al. 2005) with the C-terminal helix extending into that of the PH domain. The PH domain is a beta sandwich flanked by an amphipathic helix. Well-conserved helices in the DH domain bind directly to the switch regions of the GTPase to allow for catalytic activity (Fig. S2). Currently, there are at least 78 structures in the Protein Data Bank from X-ray crystallography, cryo-electron microscopy, and nuclear magnetic resonance that contain a Dbl RhoGEF DH domain (Ravala and Tesmer 2024). Of the publications associated with these structures, only about half explicitly include freezing as part of their standard purified protein storage protocol, and the rest are ambiguous on storage conditions. These proteins are routinely frozen with 5-10% glycerol or without cryoprotectant, yet there have not been systematic reports on either the consequences of freezing these proteins or the possible effects of cryoprotective agents (CPAs) on their activities, potentially impacting rigor and reproducibility in the field.

In addition to their normal biological function, RhoGEFs are implicated in driving a variety of diseases, including cancer, diabetes, hypertension, and neurodegenerative diseases (Blangy 2018; Fine et al. 2009). Guanine nucleotide exchange activity assays (Blangy 2018; Blaise et al. 2021; Ravala et al. 2024) and thermostability measurements (Cash et al. 2020) employing purified proteins are commonly used to characterize disease-relevant RhoGEF variants and evaluate potential chemical inhibitors, informing our understanding of how RhoGEFs can propel disease. Before deriving conclusions from these types of *in vitro* experiments, it is essential to ensure that the involved protein samples are well-behaved and functionally intact. Unless these samples are exclusively freshly prepared, understanding how freezing and storage conditions affect the activity and stability of a specific RhoGEF protein is critical for interpreting experimental results and ensuring reliable biochemical characterization of that RhoGEF. Cryopreservation of proteins through flash freezing in liquid nitrogen (lN_2_) and subsequent storage at ultralow temperatures is common practice to preserve biological materials. However, physical consequences of ice formation, like cold-induced unfolding and cryoconcentration, have been shown to disrupt protein function (Singh 2019). While rapid freezing of enzymes with lN_2_ has been widely recommended for preservation of activity (Nema and Kenneth 1993; Shikama and Yamazaki 1961), some proteins are damaged in this process (Strambini and Gabellieri 1996; Cao et al. 2003). CPAs like glycerol (Polge et al. 1949) and sucrose are broadly effective at mitigating these effects, but still some proteins remain sensitive to damage despite the addition of CPA.

Based on anecdotal evidence, our lab noticed that purified RhoGEF proteins may be sensitive to freezing and wanted to further probe this observation. We selected three fairly well-studied RhoGEFs, P-Rex1, P-Rex2, and PDZ-RhoGEF (PRG), and purified their tandem DH/PH domains. We then examined how their activities and thermostabilities change without and with different CPAs, both before and after freezing. These proteins differ in GTPase specificity and biochemical characteristics, allowing us to explore whether CPA and freezing effects may be generalizable across the Dbl RhoGEF family. Our results reveal that freezing alters activity measurements unpredictably within each GEF protein sample and, furthermore, even the addition of CPA to samples, without freezing, increases the spread of activity measurements collected and can artificially increase activity. Furthermore, we demonstrate that, at least with P-Rex2 DH/PH, changes in activity and thermostability from freezing or addition of CPA are not reflected as conformational changes that can be detected with size-exclusion chromatography coupled with small-angle X-ray scattering (SEC-SAXS). The underlying mechanisms responsible for freezing- and/or cryoprotectant-induced changes to GEF function remain unknown, but these findings underscore that these factors can unpredictably influence RhoGEF function. Thus, assays using frozen proteins without accompanying characterization of the effects of freezing and CPA on those proteins should be interpreted with caution.

## Results

### Insights into Dbl RhoGEF DH/PH biochemical differences that could potentially confer differences in stability

The Dbl RhoGEF family of proteins are characterized by a DH catalytic domain that binds GTPases and is almost always followed by a PH domain (Rossman et al. 2005). Despite this conserved architecture, the biochemical characteristics of this conserved domain tandem can be divergent, as RhoGEFs exhibit varying specificities for different GTPases. To investigate how biochemically disparate RhoGEF DH/PH domains may be, we chose four commonly studied RhoGEF proteins, P-Rex1, P-Rex2, PDZ-RhoGEF (PRG), and p63RhoGEF and one RhoGEF for which there is very little biochemical information available, ARHGEF39. P-Rex1 and P-Rex2 are two closely related family members, each sharing the same overall domain architecture and specificity for Rac GTPases (Jones and Ellisdon 2024). PRG and p63RhoGEF are G protein-regulated family members specific for RhoA (Aittaleb et al. 2010). ARHGEF39 is unique within the family, as the expressed protein consists of only the DH/PH domain tandem without any additional accessory domains, suggesting that it could represent a relatively stable isolated DH/PH. It is reportedly specific for RhoA (Anijs et al. 2022), but this protein is vastly understudied. For our comparison of these five proteins, we analyzed their DH/PH structures and corresponding electrostatic surface potentials, calculated their isoelectric points (pI), and constructed a structure-based sequence alignment (Fig. S1&S2). AlphaFold3 (Abramson et al. 2024) was used to predict the structure of the DH/PH tandem for P-Rex2 and ARHGEF39, as there are currently no published structures of the DH/PH tandems of these proteins.

Although the overall structures of these DH/PH domains are very similar, they exhibit some notable differences (Fig. S1&S2). In comparing sequence identity, the closely related P-Rex1 and P-Rex2 are 71% identical, but the identities of the other DH/PH domains to one another is only around 20%, even for RhoGEFs that share RhoA specificity (Fig. S2). In comparing surface electrostatics, most of the five DH domains are very similar in the character of their GTPase-binding sites apart from ARHGEF39, which is less electropositive in nature (Fig. S1; left column of right panel). In contrast, on the surface of the DH/PH domains opposite the GTPase-binding site, there is much more variability in electrostatic surface potential (Fig. S1; right column of right panel). Additionally, while the isoelectric points of P-Rex1, P-Rex2, and PRG are very similar at a calculated 8.5-8.7, the pI of p63RhoGEF and ARHGEF39 are much different at 6.7 and 9.8, respectively. This has important implications for working with the isolated DH/PH domains, as a purified protein is less stable in a buffer with a pH very close to the pI of the protein, suggesting that a purification buffer that works well for one of these proteins may destabilize another one. Collectively, this suggests that even though isolated RhoGEF DH/PH tandem domains are structurally similar overall, they can exhibit biochemical differences that might impact their behavior and stability in purified form.

### Glycerol and sucrose increase variation in activity measurements of freshly purified RhoGEF DH/PH proteins

Due to the biochemical similarity of P-Rex1 and P-Rex2 DH/PH domains and the divergence of PRG, we selected these three for analysis. We also chose to use isolated DH/PH tandems rather than full-length RhoGEFs to examine effects on only the catalytic core of the GEF and to limit confounding variables like intramolecular interactions from additional regulatory domains. Working with the isolated DH/PH also offers the technical advantage of utilizing bacterial expression, enabling production of large amounts of protein which allows for longitudinal analyses of biological replicates. In our hands, we observed that the activities of these proteins change after freeze-thaw.

First, we wanted to establish any broad effects on these proteins from the addition of cryoprotectant agents (CPAs) and determine if CPAs affect baseline activity measurements of freshly purified P-Rex1, P-Rex2, and PRG DH/PH tandems. Glycerol and sucrose are commonly used, powerful CPAs which promote folding and protein stability through preferential exclusion of the CPA from the protein hydration-shell, improving preservation of protein function (Lee and Timasheff 1981; Bolen and Baskakov 2001; Hirai et al. 2018; Simončič and Lukšič 2021). Thus, we analyzed protein activity of samples prepared without cryoprotectant, with 5-20% glycerol, or with 5-10% sucrose immediately after purification.

To test the activity of each RhoGEF DH/PH tandem in each CPA condition, we used a GEF activity assay which reports the increase in association between a soluble GTPase and a fluorescent, non-hydrolyzable GTP analogue (mant-GTP) (Fig. 1). For each biological replicate, activity was normalized to the cryoprotectant-free condition. P-Rex1 shows no significant changes in activity with respect to CPA condition, but the 95% confidence interval (CI) strongly increases across all conditions (Fig. 1A), suggesting the presence of CPA increases variability between replicates and makes data interpretation more difficult. P-Rex2 exhibits increased activity in 20% glycerol and 5% sucrose conditions as well as a strong increase in CI across all CPA conditions (Fig. 1B). PRG shows increased activity with 10% glycerol and both sucrose conditions (Fig. 1C). PRG presents an increased CI in most CPA conditions except 5% sucrose, although it is unclear why the data are much tighter in this condition as compared to the other conditions or samples. Together, these data indicate that the addition of glycerol or sucrose broadly increases variation in the measured activities of the proteins tested. However, these conditions do not uniformly affect activity across different proteins with any distinguishable patterns, even among closely related RhoGEFs.

**Figure 1.**
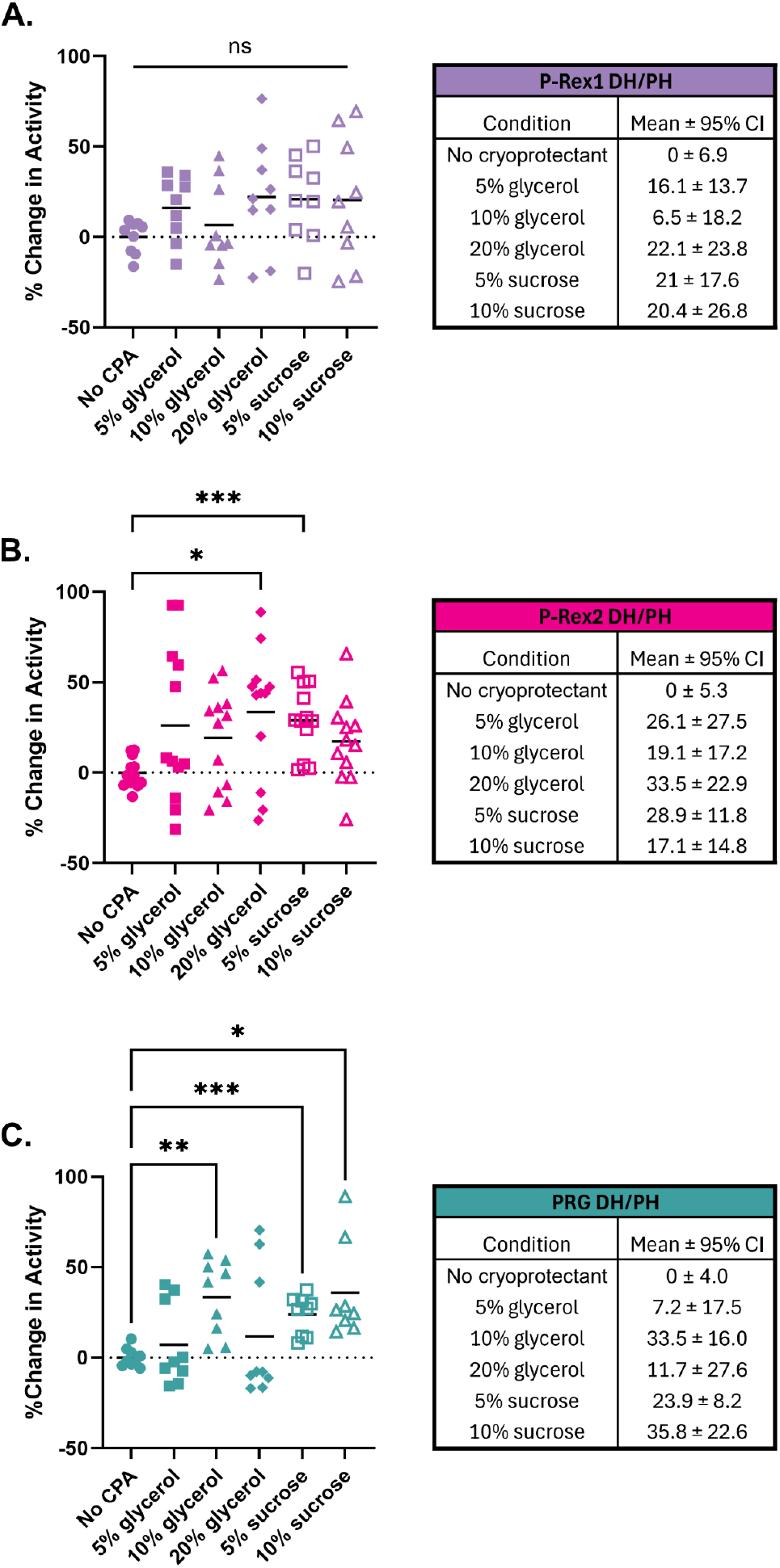
In unfrozen samples, glycerol and sucrose generally increase the variability of activity measurements of isolated RhoGEF DH/PH domain tandems. Following purification, all isolated DH/PH tandems were prepared in different cryoprotectant conditions. One hundred nM P-Rex1 DH/PH (A) and 200 nM P-Rex2 DH/PH (B) were assessed against Rac1, while 100 nM PRG DH/PH (C) was assessed against RhoA. For each biological replicate, activity was normalized to the activity of the no cryoprotectant sample to evaluate the changes in activity solely from the addition of cryoprotectant. Statistical significance was determined using Brown-Foresythe and Welch ANOVA test with a Dunnett’s T3 multiple comparisons test with individual variances computed for each comparison. ns = not significant, * p < 0.05, ** p < 0.01, *** p < 0.005.

### Isolated DH/PH protein activity is widely affected by freezing

It is common practice to freeze purified proteins and store them for later use, providing convenience over purifying them fresh for each experiment. However, some purified enzymes have been reported to lose some or all of their activity during the freeze-thaw process, including α-amylase (Whittam and Rosano 1973), catalase (Shikama and Yamazaki 1961), phosphofructokinase (Carpenter et al. 1987), and lactate dehydrogenase (Carpenter and Crowe 1988). To assess how RhoGEFs may be affected, we measured the activities of the isolated RhoGEF DH/PH tandems in each CPA condition after flash freezing and storage at -80°C for varying periods of time. We also assessed these samples by SDS-PAGE. After up to six months, we found no significant changes to the proteins as visualized by SDS-PAGE, indicating that there is no detectable degradation or irreversible aggregation of these samples upon freezing and storage (Fig. S3).

After one month, the activity of frozen P-Rex1 DH/PH without CPA and low levels of glycerol or sucrose decreases (Fig. 2). However, higher concentrations of glycerol (10-20%) and 10% sucrose seem to mitigate this effect. By the six-month timepoint, only the 10% glycerol sample consistently maintained comparable levels of P-Rex1 DH/PH activity to the fresh sample. We also observed an unexpected phenomenon where, in conditions with low levels of CPA, after an initial drop, P-Rex1 activity rises back to levels indistinguishable from the fresh sample at the six-month timepoint.

**Figure 2.**
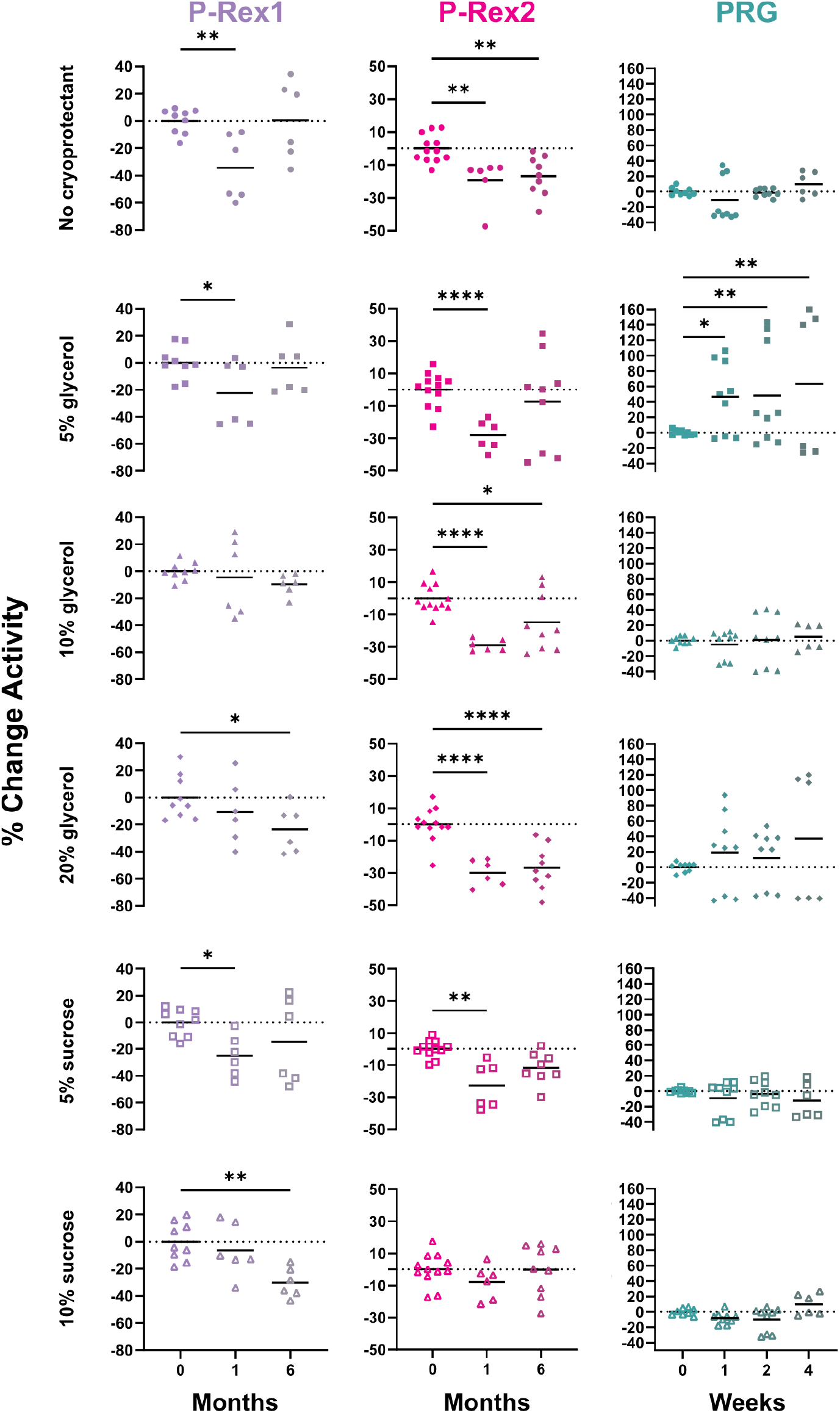
Isolated RhoGEF DH/PH activity changes over time and/or becomes less reproducible after freezing. One hundred nM P-Rex1 and 200 nM P-Rex2 DH/PH GEF activity was assessed against Rac1 while 100 nM PRG DH/PH GEF activity was assessed against RhoA. Each protein under each CPA condition was tested over various time points after freezing. Each biological replicate was normalized to the activity of the freshly purified protein. Ninety-five percent confidence intervals (CI) of these data are shown in Fig. S4. Statistical significance was determined using a two-way ANOVA test with a post hoc Dunnett’s test for multiple comparisons. * p < 0.05, ** p < 0.01, **** p < 0.001

Compared to P-Rex1, P-Rex2 DH/PH is much less resilient to freezing in most of the tested conditions. After one month, P-Rex2 DH/PH activity decreased in all storage conditions except 10% sucrose (Fig 2). After six months, the 10% sucrose sample maintained activity levels comparable to the fresh protein sample. Interestingly, we observed that the activity of P-Rex2 DH/PH stored in low CPA concentration conditions returned to near-fresh activity levels at six months, similar to P-Rex1. While the differences in the mean activities of these samples are statistically insignificant, the spread of these data are wider than that of the fresh sample as shown in the 95% confidence intervals (CI) of the data (Fig. S4).

PRG DH/PH activity was tested over a shorter period than the P-Rex proteins. Despite this, PRG revealed a wide spread between data (Fig. S4). While PRG DH/PH activity is preserved in most conditions within the 4-week time frame, there are strange trends within these data, making it unlike either P-Rex protein and difficult to interpret. For example, PRG DH/PH stored in 5% glycerol shows opposite trends between biological replicates where one shows a dramatic increase in activity after freezing while others decrease in activity. The 20% glycerol condition shows a similar occurrence.

Altogether, these data show that freezing affects each GEF in different ways with no clear or generalizable trends. Both P-Rex DH/PH samples are affected by freezing, generally manifested as decreases in activity, but surprisingly do not have the same responses across the different CPA sample conditions. PRG DH/PH, on the other hand, does not seem to be as greatly affected by freezing within a one-month timeframe. However, PRG shows even greater changes in variability of activity measurements than either P-Rex sample, and perturbations from freezing in certain glycerol conditions lead to increases in activity, unlike P-Rex samples. Overall, RhoGEF DH/PH activity measurements seem to be widely impacted by freezing.

### Thermostability of DH/PH proteins generally does not change at early time points post-freezing

Next, we wanted to assess the protein quality and stability at each of these time points to help explain the changes and variation in activity measurements. We characterized how each storage condition affects RhoGEF thermostability over the time course tested using differential scanning fluorimetry (DSF). To our surprise, protein thermostability did not correspond to observed changes in activity. Prior to freezing, the melting temperature of freshly purified P-Rex1 DH/PH across conditions with and without CPA remained consistent at ∼38°C and was preserved after one month of storage (Fig. 3). At the six-month time point, there is a significant decrease in thermostability across all conditions along with an increase in variability between measurements. With unfrozen P-Rex2 DH/PH, melting temperature increases with 20% glycerol from 41°C to 43°C but remains consistent within sucrose-containing conditions at ∼41°C (Fig. 3). Around the six-month time point post-freezing, P-Rex2 DH/PH thermostability in the no CPA condition decreases. Higher concentrations of glycerol seem to exacerbate the loss of thermostability, decreasing melting temperature by 2-3°C by six months. Unfrozen PRG DH/PH exhibits a melting temperature of ∼35°C, and, similar to P-Rex2, this increases in the presence of 20% glycerol to ∼36.5°C. The thermostability of PRG DH/PH is relatively unaffected by freezing in most conditions over the 4-week time frame with the exceptions of the 20% glycerol condition where melting temperature dropped within one week and both sucrose conditions where melting temperatures dropped after four weeks of storage.

**Figure 3.**
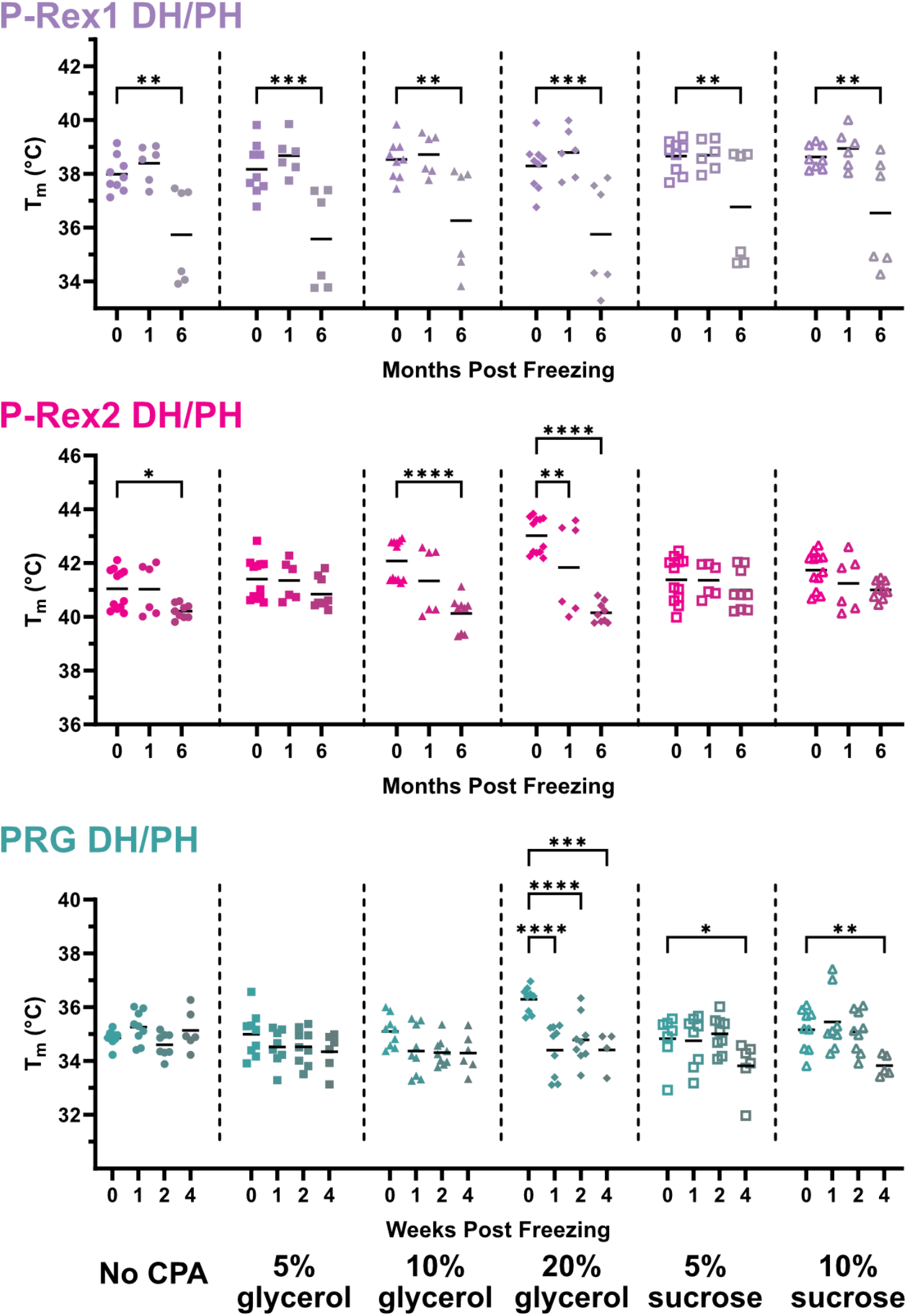
Thermostability of DH/PH tandems begins to drop at late time points. P-Rex1 DH/PH, P-Rex2 DH/PH, and PRG DH/PH thermostability was tested with differential scanning fluorimetry (DSF) across cryoprotectant conditions at various time points. Statistical significance was determined using a two-way ANOVA test with a post hoc Dunnett’s test for multiple comparisons. * p < 0.05, ** p < 0.01, *** p < 0.005, **** p < 0.001

### Long-term freezing does not affect the global structure of P-Rex2 DH/PH

Changes to thermostability and activity after storage at -80°C led us to investigate whether or not the global structure of the DH/PH tandem itself was changing, leading to these differences. To do this, we utilized size exclusion chromatography coupled with small-angle X-ray scattering (SEC-SAXS). Because P-Rex2 DH/PH activity and thermostability were affected upon freezing without CPA but preserved in 5% sucrose, we chose these two conditions and let the samples mature at -80°C for 6 months before assessing them using SEC-SAXS. We compared these samples to data previously collected by our lab on a freshly purified P-Rex2 DH/PH sample (Anderson et al. 2026; SASDB#). Both the CPA-free and the 5% sucrose samples eluted in a single symmetric peak (Fig. S5A). There is a slight delay in the elution of the frozen CPA-free sample and furthermore in the 5% sucrose sample, but this could be due to differences in column interaction. In both samples, there are no signs of aggregation or repulsion at low q in either the scattering intensity plots (Fig. 4A-B) or Guinier analyses (Fig. S5B-C). Dimensionless Kratky analysis also confirmed that, after freezing and storage, both samples maintain a compact and folded structure with well-defined peaks corresponding to globular proteins and near-identical profiles to the fresh P-Rex2 DH/PH (Fig. 4C). The molecular weight calculated for each sample corresponds to that of a P-Rex2 DH/PH monomer expected to be ∼44 kD (Table 1). Interestingly, the correlation volume of fresh P-Rex2 DH/PH was reported at 41.6 kD, but the frozen CPA-free and 5% sucrose samples were 42.5 and 39.3 kD, respectively. Given the comparable radius of gyration (Rg) between these samples (Table 1), it is unlikely that the difference in molecular weight is significant. Finally, the pair distance distribution function, P(r), indicated D_max_ values of 81.6 Å (CPA-free) and 80.1 Å (5% sucrose) (Fig. 4D, Table 1), again demonstrating that there are not any significant global differences in conformation, shape, or oligomeric state. Therefore, the changes that lead to decreases in thermostability and activity are not necessarily related to large scale conformational changes and may be due to other sources of damage.

**Table 1.**
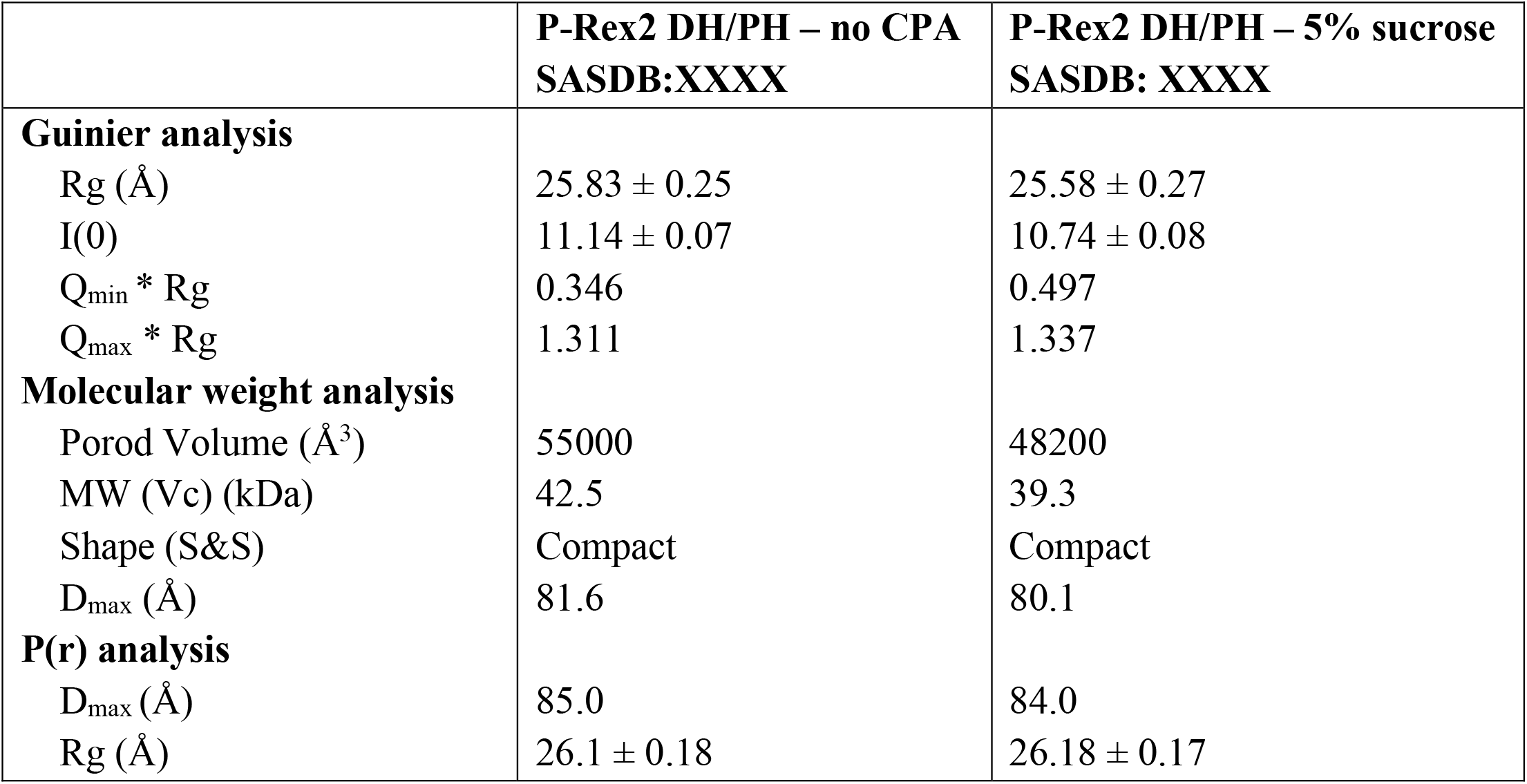
SEC-SAXS data analyses.

**Figure 4.**
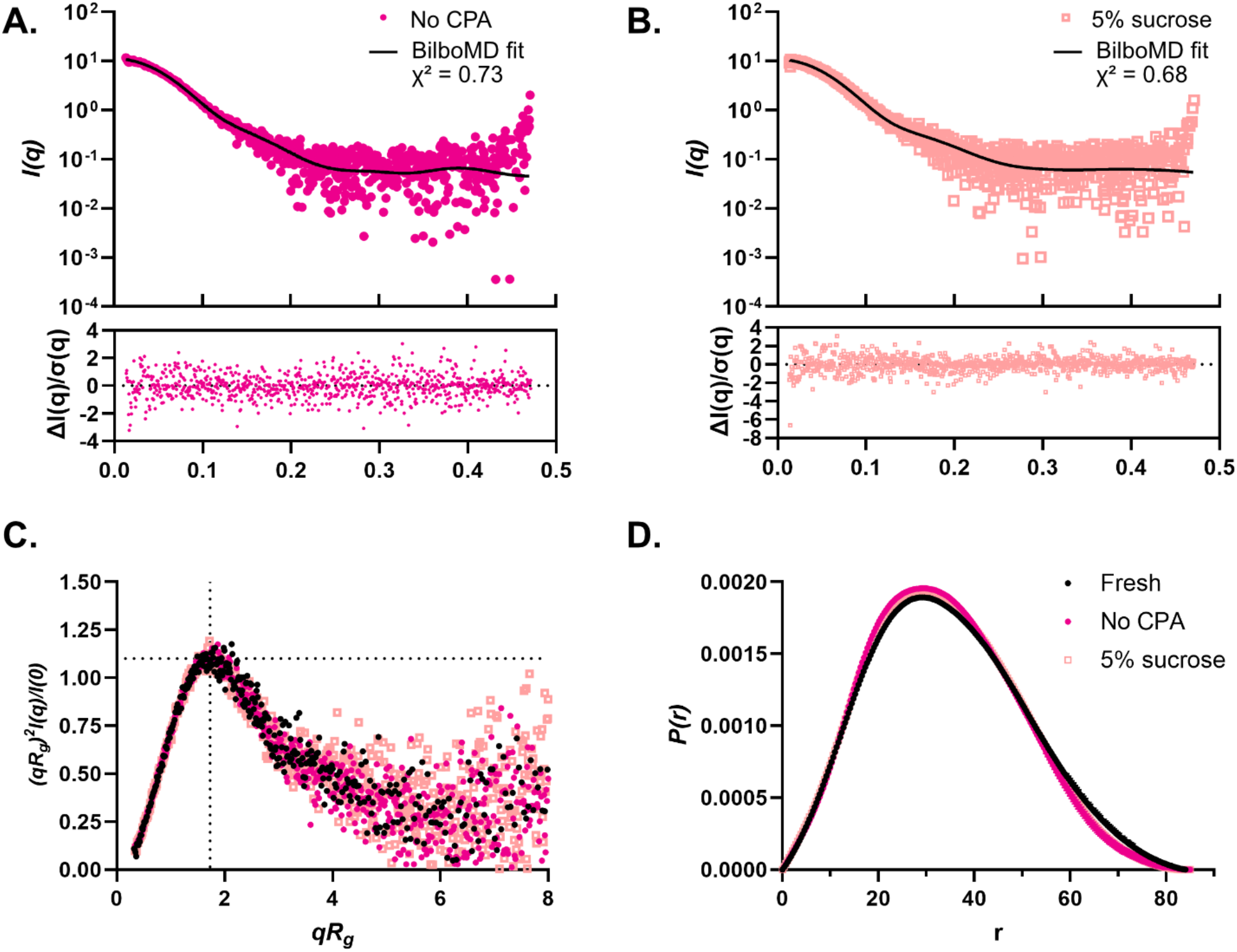
SEC-SAXS of frozen P-Rex2 DH/PH does not reveal large changes to protein conformation. Scattering intensity plots of P-Rex2 DH/PH after freezing in (A) no CPA and (B) 5% sucrose after 6 months of storage. C. Dimensionless Kratky plot comparing freshly purified P-Rex2 DH/PH (black circles) and frozen P-Rex2 DH/PH samples with no CPA (magenta circles) or 5% sucrose (peach squares). There are no significant differences in shape between the fresh or frozen samples. D. Pair distance distribution function of freshly purified P-Rex2 DH/PH and samples frozen without CPA or 5% sucrose.

## Discussion

In this study, we examined how flash freezing and storage at ultralow temperatures affect the RhoGEF activity and thermostability of isolated DH/PH tandems from P-Rex1, P-Rex2, and PRG. We also evaluated whether or not glycerol or sucrose could preserve these properties during storage. The answers to these questions are complicated: each isolated DH/PH tandem responded to different freezing and storage conditions in unique ways. Even closely related RhoGEFs, P-Rex1 and P-Rex2, did not produce the same response to the same CPA conditions. One of the few widely generalizable trends we observed is that both the addition of cryoprotectants and prolonged storage of these purified RhoGEFs increases variability and reduces reproducibility of activity measurements. Compounding this phenomenon, the mant-GTP exchange assay itself presents technical challenges that can further complicate data interpretation. Perturbations in how the assay is performed, including but not limited to the order in which reagents are mixed, how quickly the initiated reaction is measured, and selection of RhoGEF concentration in the assay, can lead to variable results. Because of this, we spent substantial time optimizing how we perform the assay to make it as consistent as possible, as evidenced by low variability within unfrozen samples, especially those without CPA. Even after optimization of this workflow and attentiveness to consistent handling techniques, adding CPA to or freezing RhoGEF samples make the resulting data more difficult to interpret. Moreover, the amount of cryoprotectant carried into the activity assay is minimal. Even at the highest glycerol concentration tested, the final concentration added to the reaction is less than 0.5%. Thus, it is unlikely that cryoprotectants directly affect the assay itself but rather influence the apparent biochemical qualities of the proteins.

We also found that a loss of activity and/or increase in variability does not necessarily correspond to a decrease in thermostability. Moreover, despite clear changes in activity and thermostability in P-Rex2 DH/PH, SEC-SAXS analysis revealed no detectable changes in overall conformation or flexibility compared to fresh protein. Given the limitations of SEC-SAXS for resolving structural changes as a low-resolution method, this suggests that more subtle shifts in regulatory or dynamic states may underlie the observed effects, requiring more high-resolution solution-based methods like hydrogen-deuterium exchange mass spectrometry to determine what causes these changes. While the results were not definitive, the absence of clear trends among GEFs and cryoprotectants in preserving activity and thermostability serves as an important reminder that protein handling conditions must be carefully characterized to ensure reproducibility.

While there are few generalizable trends between GEFs, these data indicate potential appropriate frozen storage conditions for individual RhoGEFs. For example, P-Rex1 DH/PH stored in 10% glycerol appears to retain catalytic activity for up to six months despite a noticeable reduction in thermostability over the same period. This suggests that glycerol may help preserve function even with subtle structural destabilization. For P-Rex2 DH/PH, 10% sucrose may serve as a suitable cryoprotectant under the conditions tested. Furthermore, these CPA concentrations could represent suitable storage conditions for full-length P-Rex proteins as well. However, these practices need to be validated with equally rigorous work with the full-length proteins before adopting them into protocols. In contrast, PRG maintained both activity and thermostability for at least one month while frozen under CPA-free conditions.

Together, these findings caution against assuming that frozen RhoGEFs retain their functional properties and emphasize that biochemical assays using frozen protein should be interpreted carefully. Moreover, these findings complicate direct comparisons between studies using frozen proteins, especially those from different labs where published information on protein handling and storage is not clear or provided. Our study calls attention to potential alarming issues around reproducibility in the field of biochemically characterizing purified RhoGEF proteins. More broadly, this work serves as a cautionary reminder that not all proteins– regardless of how well-conserved or similar in function– are affected the same by freezing, even under the same conditions.

## Materials and Methods

### Cloning and Plasmids

P-Rex1 DH/PH, P-Rex2 DH/PH, and soluble Rac1 expression constructs were described previously (Cash et al., 2016; Anderson et al., BioRxiv; Kristelly et al., 2004). pDONR221 PRG was obtained courtesy of Center for Personalized Diagnostics through DNASU (HsCD00860223) (Seiler et al. 2014). PRG DH/PH (residues 729-1081) were cloned into a pMALc2H_10_T vector with a Tobacco Etch Virus (TEV)-cleavable N-terminal maltose binding protein (MBP)-10xHis tag. pLV_DORA_RhoA_IRES was generated by the Hayer lab and gifted to us through Dr. Sean Collins (Marshall-Burghardt et al. 2024) Soluble RhoA (residues 1-189, excluding the CAAX site) was cloned into the same pMALc2H_10_T vector with a TEV-cleavable N-terminal MBP-10xHis tag.

### Protein production, purification, and handling

Soluble Rac1 and RhoA were expressed in Rosetta (DE3) pLysS *E. coli* cells (Invitrogen, USA) and purified as previously described (Cash et al. 2016).

P-Rex and PRG DH/PH constructs also were expressed in Rosetta (DE3) pLysS *E. coli* cells as N-terminal His-tagged MBP fusion proteins. Cells were grown in Terrific Broth to an OD_600_ of 0.8, induced with 0.1 mM IPTG at 16-20 °C, harvested after 16-20 hours, and then flash frozen and stored at -80 °C. Cell pellets were thawed and resuspended in 9 ml 20 mM HEPES (pH 8), 300 mM NaCl, 0.1 mM EDTA, 2 mM DTT and protease inhibitor cocktail (125 nM aprotinin, 6.8 µM leupeptin, 0.15 nM soybean trypsin inhibitor, 1 µM E-64, and 1 µM bestatin) per 1 g cell pellet. Cells were then homogenized with a dounce and lysed with an Avestin EmulsiFlex-C5 High Pressure Homogenizer (Avestin, Inc.; Ottawa, ON). Lysate was clarified by ultracentrifugation at 40,000 rpm for 45 min in a Type 45 Ti rotor (Beckman Coulter; Brea, CA). The supernatants were then filtered through a glass fiber filter.

P-Rex DH/PH clarified supernatant was incubated with Ni-NTA resin for 30-60 min. Resin was washed with buffer A [20 mM HEPES (pH 8), 300 mM NaCl, 2 mM DTT] followed by buffer A containing 10 mM imidazole, and proteins were eluted with buffer A containing 250 mM imidazole. Elutions were then simultaneously dialyzed into buffer containing 20 mM HEPES (pH 7), 200 mM NaCl, 2 mM DTT, and 10% glycerol and treated with TEV protease overnight to remove the N-terminal His-tagged MBP at a 1:2 molar ratio of TEV:MBP-P-Rex DH/PH. The cleaved MBP-His was then captured by an additional pass over Ni-NTA resin. The flow through was then processed over a HiTrap SP Sepharose Fast Flow column (Cytiva; Marlborough, MA) in 20 mM HEPES (pH 7) and 2 mM DTT and eluted over an NaCl gradient from 0-0.5 M over 4 CV. Fractions containing P-Rex DH/PH were pooled and concentrated to ∼2 mg/ml, diluted to 1 mg/ml with buffer [20 mM HEPES (pH 7), 100 mM NaCl, and 2 mM DTT] containing 0-20% glycerol or 0-10% sucrose, and thoroughly and gently pipet mixed. Fresh protein was promptly used for assays where DSF was conducted within 15 minutes and the GEF activity assay was performed within 30-90 minutes of sample assembly. The protein was then immediately flash frozen in lN_2_ and stored at -80 °C in 60 uL aliquots. Frozen P-Rex1 DH/PH samples were thawed on ice and then thoroughly pipet mixed. Frozen P-Rex2 DH/PH samples were thawed under cold running water, thoroughly pipet mixed, and then placed on ice.

PRG clarified supernatant was incubated with amylose resin for 1 hour. Resin was washed with buffer A and eluted with buffer A containing 10 mM maltose. Elutions were then simultaneously dialyzed into buffer containing 20 mM HEPES (pH 8), 100 mM NaCl, and 2 mM DTT and treated with TEV protease overnight to remove the N-terminal His-tagged MBP at a 1:6 molar ratio of TEV:MBP-PRG DH/PH. The dialyzed sample was then incubated with Ni-NTA resin for 1 hour to capture the cleaved MBP-His. The flow through was then processed over ENrich SEC 70 (BioRad; Hercules, CA) in 20 mM HEPES (pH 8), 200 mM NaCl, 2 mM DTT. Fractions containing PRG DH/PH were pooled and concentrated to ∼2-5 mg/ml and diluted to 1 mg/ml with buffer [20 mM HEPES (pH 8), 200 mM NaCl, 2 mM DTT] containing 0-20% glycerol or 0-10% sucrose. Fresh protein was promptly used for assays where DSF was conducted within 15 minutes and the GEF activity assay was performed within 30-90 minutes of sample assembly. The protein was then immediately flash frozen in lN_2_ and stored at -80 °C in 82 uL aliquots. Frozen PRG DH/PH samples were thawed at 4 °C, thoroughly pipet mixed, and then placed on ice.

### GTPase Exchange Activity Assay

A FRET-based guanine nucleotide exchange assay was used to assess the activity of the fresh and frozen isolated DH/PH protein samples. P-Rex DH/PH samples were diluted from 1 mg/ml (∼22.7 µM) to 0.1 mg/ml with assay buffer [20 mM HEPES (pH 8), 100 mM NaCl, 5 mM MgCl_2_, 1 mM DTT, and 5% glycerol]. PRG DH/PH samples were diluted from 1 mg/ml (24.15 µM) to 0.06 mg/ml (1.625uM) with assay buffer. A final concentration of 100 nM P-Rex1 or 200 nM P-Rex2 DH/PH was added to 2 µM Rac1 or 100 nM PRG DH/PH was added to 2 µM RhoA in assay buffer. The reaction was then initiated by adding 0.8 µM 2’/3’-O-(N-methyl-anthraniloyl)-guanosine-5’-diphosphate (mant-GTP), a non-hydrolyzable form of GTP. The reactions were excited at 280 nm and the fluorescence intensity at 450 nm was measured in 30s intervals on a SpectraMax M5 (Molecular Devices; San Jose, CA) plate reader for a total of 15 min. Fluorescence curves were fit to a one-phase association model using GraphPad Prism (San Diego, CA). The k value obtained from the one-phase association model was used to represent GEF activity. A range of DH/PH protein concentrations were empirically tested to determine the most appropriate one to use. Because of instrumentation limitations which restricted obtaining initial rates in this assay, lower DH/PH concentrations that strongly fit to a one-phase association equation were favored. In all RhoGEF activity assays, each biological replicate was normalized within itself. For the analysis of CPAs on freshly purified RhoGEF activity (Fig. 1), the freshly purified CPA-free sample was used for normalization within each biological replicate. For the analysis of freezing in each CPA condition (Fig. 2), the unfrozen purified protein in each CPA condition was used for normalization within each biological replicate. We selected this strategy to analyze the trends in RhoGEF activity from CPA-addition or freezing and to minimize the effect of slight variation between biological replicates in data interpretation.

### Differential Scanning Fluorimetry

Experiments were performed on a CFX96 Real-Time PCR System (BioRad; Hercules, CA). Purified P-Rex DH/PH was diluted to a final concentration of 0.5 mg/ml in a buffer containing 20 mM HEPES (pH 8), 100 mM NaCl, and 2 mM DTT. Purified PRG DH/PH was diluted to a final concentration of 0.6 mg/ml in a buffer containing 20 mM HEPES (pH 8), 200 mM NaCl, and 2 mM DTT. 10X Sypro or Prolite (AAT Bioquest; Pleasanton, CA) Orange dye was added to each condition, and fluorescence was monitored across temperatures from 22 °C to 95 °C, incrementing by 1 °C per minute. Fluorescence signal was measured in the dedicated FRET channel as per manufacturer specifications. The protein melting temperature (T_m_) was determined by fitting the fluorescence data to a Boltzmann sigmoidal curve and calculating the inflection point in GraphPad Prism.

### Size exclusion chromatography coupled to small angle x-ray scattering (SEC-SAXS)

Purified P-Rex2 DH/PH was frozen without cryoprotectant or with 5% sucrose and stored at -80 °C for six months. Samples were thawed under cold water, gently pipet mixed, and then thoroughly buffer exchanged into 20 mM HEPES (pH 7), 200 mM NaCl, 2 mM DTT, and 1% glycerol in a 10 kD Amicon (MilliporeSigma; Burlington, MA) concentrator using 2 minute 8000 x g spins. During buffer exchange, both samples showed signs of sample precipitation. To remove aggregated protein, samples were spun for 10 minutes at 14,000 x g and decanted to separate soluble from pelleted protein. Soluble samples were shipped to the Structurally Integrated Biology for Life Sciences (SIBYLS) beamline for data collection and analyzed as previously described (Anderson et al. 2026).

## Supplementary Material

Anderson_Freezing_GEFs_Supp.pdf is a pdf containing supplemental figures S1-S5.

## Data Sharing and Availability Statement

The SAXS data for frozen P-Rex2 DH/PH without cryoprotectant and with 5% sucrose can be accessed as SASDB entries XXXX and XXXX, respectively.

## Acknowledgments

Research reported in this publication was supported by the National Institute of General Medical Sciences of the National Institutes of Health under Award Number R35GM146664 (to J.N.C.). This project was supported in part by Administrative Coordinating Council of Deans (ACCD) College of Biological Sciences funds received from the UC Davis Clinical and Translational Science Center, which is supported by award UL1 TR001860 from the NIH National Center for Advancing Translational Sciences (to J.N.C.). SAXS data were collected at the Advanced Light Source (ALS) at the SIBYLS beamline, a national user facility operated by Lawrence Berkeley National Laboratory on behalf of the Department of Energy, Office of Basic Energy Sciences, through the Integrated Diffraction Analysis Technologies (IDAT) program, supported by DOE Office of Biological and Environmental Research. Additional support comes from the National Institute of Health project ALS-ENABLE (P30 GM124169). We would like to thank Dr. Daniel Elnatan for suggesting sucrose as an alternative cryoprotectant agent and inspiring this study.

## Author contributions

L.K.A. conceptualized and led this study. P-Rex2 experiments were performed by L.K.A, P-Rex1 experiments were performed by H.M., and PRG experiments were performed by E.B. L.K.A. pooled and analyzed all final data and made the corresponding figures. J.N.C performed the structure and sequence analysis but mainly sat the bench on this one and fulfilled a cheerleading role.

## CRediT author contributions

Conceptualization - L.K.A and J.N.C.

Data curation - L.K.A

Formal analysis - L.K.A, E.B., and H.M.

Funding acquisition - J.N.C.

Investigation - all authors

Methodology - L.K.A and J.N.C.

Project administration - J.N.C.

Resources - J.N.C.

Supervision - L.K.A and J.N.C.

Validation - L.K.A and J.N.C.

Visualization - L.K.A and J.N.C.

Writing — original draft - L.K.A and J.N.C.

Writing — review & editing - all authors

## Abbreviations and symbols

RhoGEF: Rho guanine-nucleotide exchange factor
DH: Dbl homology
PH: pleckstrin homology
CPA: cryoprotectant agent
lN_2_: liquid nitrogen
P-Rex: phosphatidylinositol-3,4,5-trisphosphate-dependent Rac exchanger
PDZ: postsynaptic density 95, Discs large, Zonula occludens-1
PRG: PDZ-RhoGEF
SEC-SAXS: size exclusion chromatography coupled with small-angle X-ray scattering
CI: confidence interval
DSF: differential scanning fluorimetry
P(r): pair distance distribution function
TEV: Tobacco Etch Virus
MBP: maltose binding protein
mant-GTP: 2’/3’-O-(N-methyl-anthraniloyl)-guanosine-5’-diphosphate

