## Supplemental Figures for "GEF me a break: the consequences of freezing Rho guanine-nucleotide exchange factor catalytic domains"

**Contents**

Supporting Information Figures S1-S5

**Figure S1.** Comparison of structures and characteristics of DH/PH tandem domains from selected Dbl RhoGEFs.
**Figure S2.** Sequence alignment of the DH/PH structures shown in Figure S1.
**Figure S3.** RhoGEF DH/PH protein samples run on SDS-PAGE after freezing compared to fresh.
**Figure S4.** Average and spread of activity data in CPA conditions over time from Figure 2.
**Figure S5.** Frozen P-Rex2 DH/PH elution profiles and Guinier analyses.

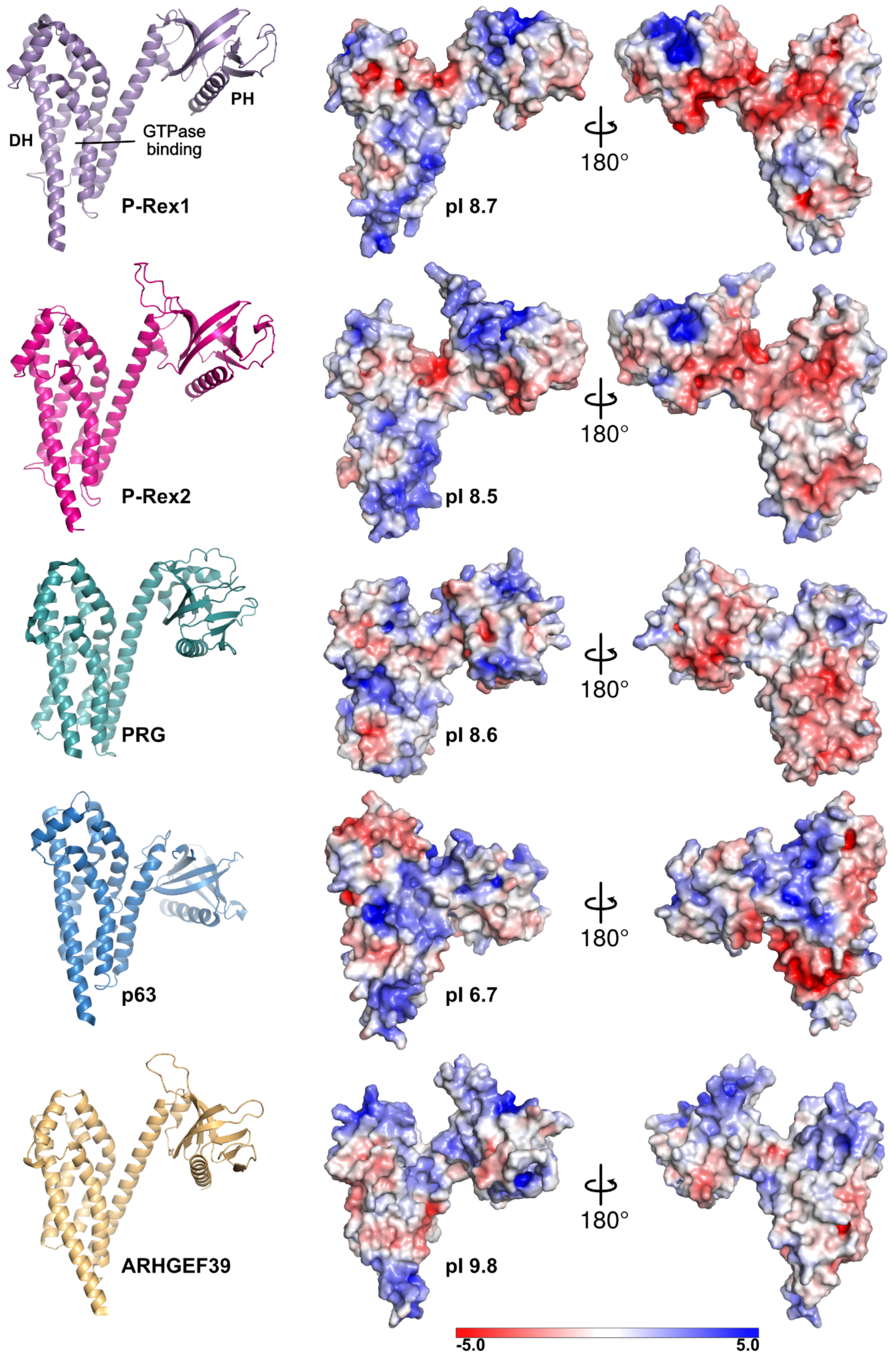


**Figure S1. Comparison of structures and characteristics of DH/PH tandem domains from select Dbl RhoGEFs.** The isolated DH/PH tandem domains of P-Rex1 (PDB ID: 5FI1), P-Rex2 (AlphaFold3 prediction), PRG (PDB ID:  IXCG), p63RhoGEF (PDB ID: 2RGN), and ARHGEF39 (AlphaFold3 prediction) are shown with structural features labeled. For those from experimental structures, only the DH/PH tandem is shown; the other components of each structure were removed. The right panel shows the electrostatic surface potential of each DH/PH calculated using Adaptive Poisson-Boltzmann Solver (APBS; Jurrus et al., 2018) electrostatics within PyMOL (Schrödinger 2010; The PyMOL Molecular Graphics System, Version 3.1.8) on a range of +/- 5.0, with blue indicating electropositive and red electronegative. In this panel, the view of the structure on the left in surface is the same view of the structure as is shown in the left panel in cartoon. Also shown for each is the isoelectric point (pI) calculated using ProtParam. The sequence used for each RhoGEF is shown in Figure S2.

```
                     α1 helix
p63       150 EEEQKKKALERSMYVLSELVET 171
PRG       731 -------REIDRQEVINELFVT 745
ARHGEF39  12  VQEQRARWERKRACTARELLET 33
PRex1     39  AARESERQLRLRLCVLNEILGT 60
PRex2     13  SAKDLEKQLRLRVCVLSELQKT 34
```

Percent identity matrix

| | | | | | |
|---|---|---|---|---|---|
| p63 | 100 | 17 | 20 | 21 | 23 |
| PRG | 19 | 100 | 17 | 20 | 19 |
| ARHGEF39 | 20 | 19 | 100 | 21 | 19 |
| PRex1 | 21 | 20 | 21 | 100 | 71 |
| PRex2 | 23 | 19 | 19 | 71 | 100 |

```
                  α1 helix        α1' helix        α2 helix
p63       EKMYVDDLGQIVEGYMATMAAQGVPE---S-LRGRDRIVFGNIQQIYEWHRDYFLQELQR 227
PRG       EASHLRTLRVLDLIFYQRMKKEN------LMPREELARLFPNLPELIEIHNSWC-EAMKK 798
ARHGEF39  ERRYQEQLGLVATYFLGILKAKG------TLRPPERQALFGSWELIYGASQELL-PYLEG 86
PRex1     ERDYVGTLRFLQSAFLHRIRQNVADSVEKGLTEENVKVLFSNIEDILEVHKDFL-AALEY 119
PRex2     ERDYVGTLEFLVSAFLHRMNQCAASKVDKNVTEETVKMLFSNIEDILAVHKEFL-KVVEE 93

                                 α3 helix                 α4 helix
p63       CLKDPD----WLAQLFIKHE-----RRLHMYVVYCQNKPKSEHVVSEFGDS--Y---F-- 271
PRG       LREEGPIIKEISDLMLARFDGPAREELQQVAAQFCSYQSIALELIKTKQRKESRFQLFMQ 858
ARHGEF39  G---------CWGQGLEGFC-----RHLELYNQFAANSERSQTTLQEQLKKNKGFRRFVR 132
PRex1     CLHPEPQSQHELGNVFLKFK-----DKFCVYEEYCSNHEKALRLLVEL-NKIPTVRAFLL 173
PRex2     CLHPEPNAQQEVGTCFLHFK-----DKFRIYDEYCSNHEKAQKLLLEL-NKIRTIRTFLL 147

          α4 helix            α5 helix                     α6 helix
p63       --EELRQQLGHRLQLNDLLIKPVQRIMKYQLLLKDFLKYYNRAGMDTADLEQAVEVMCFV 329
PRG       --EAESHPQCRRLQLRDLIISEMQRLTKYPLLLESIIKHTEGGTSEHEKLCRARDQCREI 916
ARHGEF39  --LQEGRPEFGGLQLQDLLPLPLQRLQQYENLVVALAENTGPNSPDHQQLTRAARLISET 190
PRex1     SCMLLGGRKTTDIPLEGYLLSPIQRICKYPLLLKELAKRTPGKHPDHPAVQSALQAMKTV 233
PRex2     NCMLLGGRKNTDVPLEGYLVTPIQRICKYPLILKELLKRTPRKHSDYAAVMEALQAMKAV 207

           α6 helix     αN helix                                β1 strand
p63       PKRCN----DMMTLGRLRGFEGKLTAQGK------------LLGQDTFWVTE--PEAGGL 371
PRG       LKYVNEAVKQTENRHRLEGYQKRLDATALERASNPLAAEFKSLDLTTRKMIHEGPLTWRI 976
ARHGEF39  AQRVHTIGQKQKNDQHLRRVQALLSGRQA-----------KGLTSGR-WFLRQGWLLVVP 238
PRex1     CSNINETKRQMEKLEALEQLQSHIEGWEG-----------SNLTDICTQLLLQGTL-LKI 281
PRex2     CSNINEAKRQMEKLEVLEEWQSHIEGWEG-----------SNITDTCTEMLMCGVL-LKI 255

             β2 strand   β3 strand                       β4 strand  β5 strand
p63       LSSRGRERRVFLFEQIIIFSEALGGG---------VRGGTQPGYVYKNSIKVSCLGLEGN 422
PRG       SKDKTLDLHVLLLEDLLVLLQKQDEKLLLKCHSKTAVGSSDSKQTFSPVLKLNAVLIRSV 1036
ARHGEF39  PHGEPRPRMFFLFTDVLLMAKPRPPLHLLRSGTFAC----KALYPMAQCHLSRV------ 288
PRex1     SAGNIQERAFFLFDNLLVYCKRKSRVTGSKKSTKRTKSINGSLYIFRGRINTEVMEVENV 341
PRex2     SSGNIQERVFFLFDNLLVYCKRKHRRLKNS-----KASTDGHRYLFRGRINTEVMEVENV 310

                      β6 strand    β7 strand         αC helix
p63       LQGDPCRFALTSR--------GPEGGIQRYVLQAADPAISQAWIKHVAQILESQRD 470
PRG       ATD-KRAFF---IICT-----SKLGPPQIYELVALTSSDKNTWMELLEEAVRNATR 1083
ARHGEF39  -------FGHSGGPC-GG-LLSLSFPHEKLLLMSTDQEELSRWYHSLTWAISSQKN 335
PRex1     EDG-TADYHSNGYTVTNGWKIHNTAKNKWFVCMAKTAEEKQKWLDAIIREREQRES 396
PRex2     DDG-TADFHSSGHIVVNGWKIHNTAKNKWFVCMAKTPEEKHEWFEAILKERERRKG 365
```

**Figure S2. Sequence alignment of the DH/PH structures shown in Figure S1.** Alignments were performed using Clustal Omega and then manually adjusted based on inspection of the alignment of each structure to that of P-Rex1. Percent identities are of just the sequence regions shown and calculated by Clustal Omega. Associated secondary structure is annotated based on that of P-Rex1. Loops are left unlabeled.

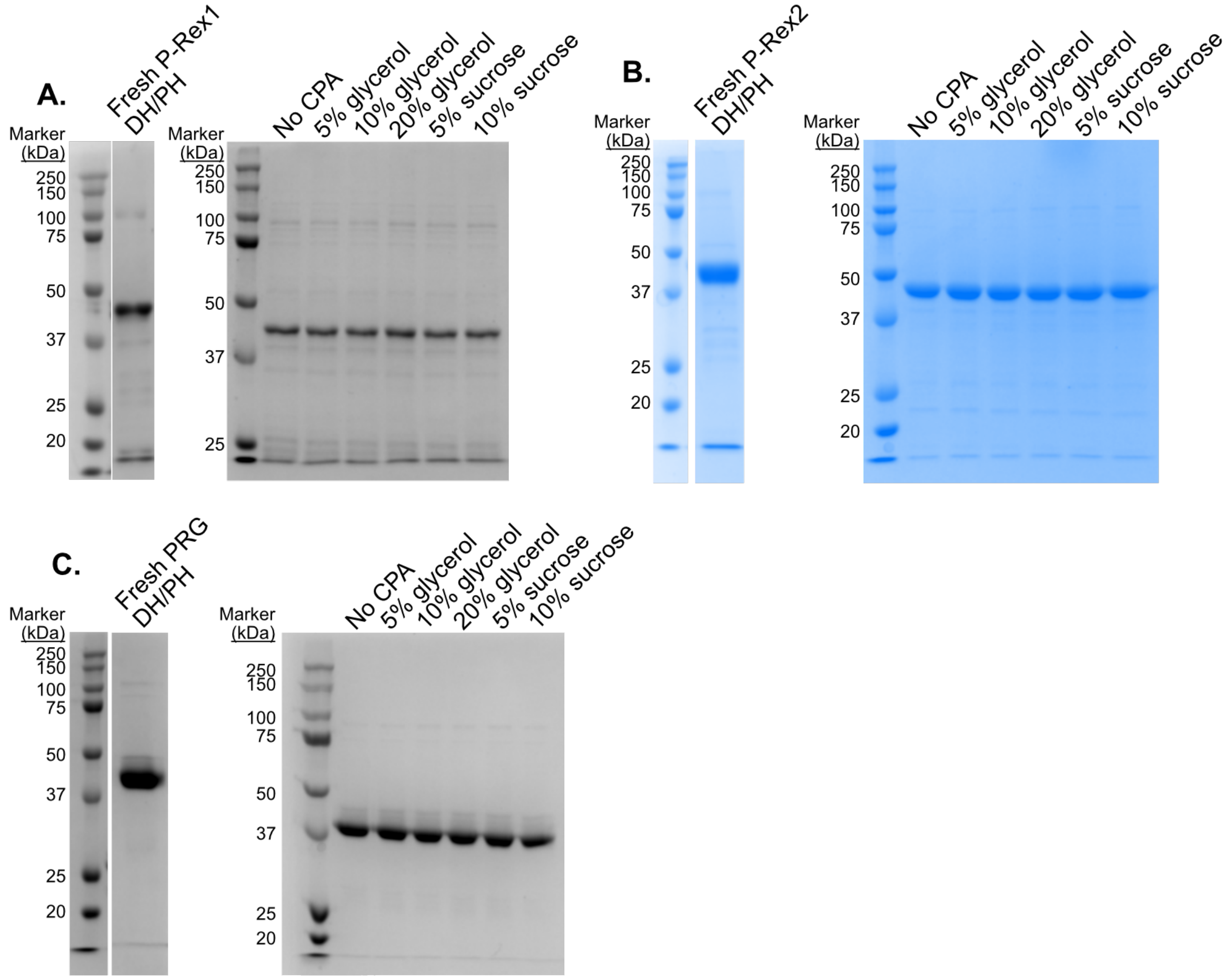


**Figure S3. RhoGEF DH/PH protein samples run on SDS-PAGE after freezing compared to fresh.** A. P-Rex1 fresh and frozen for 6 months samples. B. P-Rex2 fresh and frozen for 6 months samples. C. PRG fresh and frozen for 2 weeks samples.

| P-Rex1 DH/PH | | |
|---|---|---|
| Condition | Time Post Freezing (Months) | Mean ± 95% CI |
| No cryoprotectant | 0 | 0 ± 6.9 |
| | 1 | -34.4 ± 25.1 |
| | 6 | 0.5 ± 30 |
| 5% glycerol | 0 | 0 ± 9.3 |
| | 1 | -22.3 ± 25.2 |
| | 6 | -3.5 ± 20.8 |
| 10% glycerol | 0 | 0 ± 5.1 |
| | 1 | -4.5 ± 30 |
| | 6 | -9.7 ± 8.1 |
| 20% glycerol | 0 | 0 ± 12.5 |
| | 1 | -10.8 ± 24.9 |
| | 6 | -23.4 ± 17.8 |
| 5% sucrose | 0 | 0 ± 7.7 |
| | 1 | -25.2 ± 16.2 |
| | 6 | -14.1 ± 33.5 |
| 10% sucrose | 0 | 0 ± 10.6 |
| | 1 | -6.4 ± 20.4 |
| | 6 | -30.1 ± 11.4 |

| P-Rex2 DH/PH | | |
|---|---|---|
| Condition | Time Post Freezing (Months) | Mean ± 95% CI |
| No cryoprotectant | 0 | 0 ± 5.3 |
| | 1 | -19.4 ± 14.6 |
| | 6 | -16.7 ± 9.2 |
| 5% glycerol | 0 | 0 ± 6.7 |
| | 1 | -28.1 ± 9.5 |
| | 6 | -7.4 ± 22.4 |
| 10% glycerol | 0 | 0 ± 5.4 |
| | 1 | -29 ± 3.7 |
| | 6 | -14.8 ± 13.8 |
| 20% glycerol | 0 | 0 ± 6.6 |
| | 1 | -29.9 ± 8.4 |
| | 6 | -26.8 ± 10.4 |
| 5% sucrose | 0 | 0 ± 3.3 |
| | 1 | -22.7 ± 14.9 |
| | 6 | -11.9 ± 8.2 |
| 10% sucrose | 0 | 0 ± 6.2 |
| | 1 | -7.9 ± 11.1 |
| | 6 | -0.2 ± 11.9 |

| PRG DH/PH | | |
|---|---|---|
| Condition | Time Post Freezing (Weeks) | Mean ± 95% CI |
| No cryoprotectant | 0 | 0 ± 3.9 |
| | 1 | -10.6 ± 22.5 |
| | 2 | -1.5 ± 4.1 |
| | 4 | 9.5 ± 16.7 |
| 5% glycerol | 0 | 0 ± 2.5 |
| | 1 | 46.6 ± 35.3 |
| | 2 | 48.2 ± 50.4 |
| | 4 | 63.3 ± 99 |
| 10% glycerol | 0 | 0 ± 4 |
| | 1 | -5.2 ± 14.2 |
| | 2 | 0.8 ± 25.8 |
| | 4 | 4.7 ± 17.3 |
| 20% glycerol | 0 | 0 ± 4.5 |
| | 1 | 18.9 ± 38.8 |
| | 2 | 11.8 ± 28.5 |
| | 4 | 37 ± 89.2 |
| 5% sucrose | 0 | 0 ± 1.9 |
| | 1 | -9.2 ± 17.5 |
| | 2 | -3.9 ± 12.7 |
| | 4 | -12.4 ± 23.6 |
| 10% sucrose | 0 | 0 ± 3.7 |
| | 1 | -8.1 ± 5.8 |
| | 2 | -9.7 ± 12.2 |
| | 4 | 9.8 ± 14.5 |

**Figure S4. Average and spread of activity data in CPA conditions over time from Figure 2.**

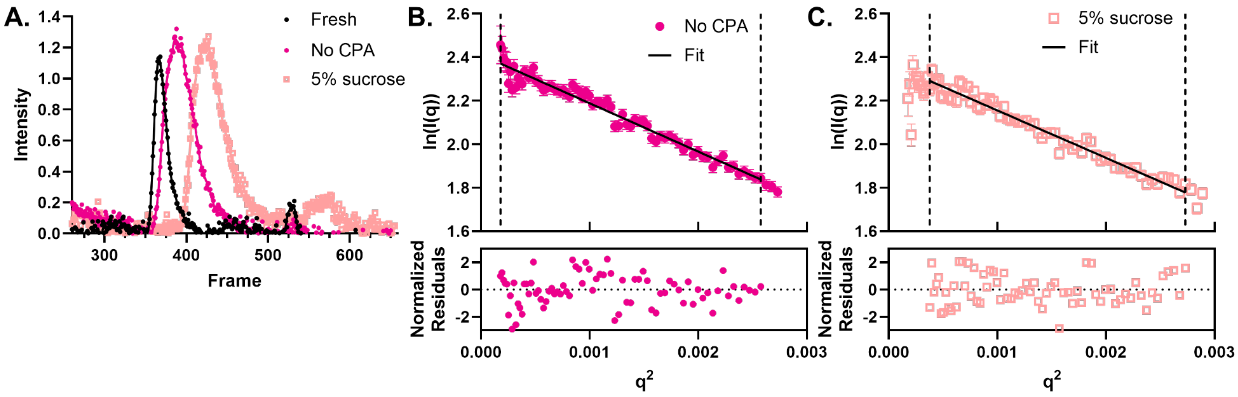


**Figure S5. Frozen P-Rex2 DH/PH elution profiles and Guinier analyses.** A. Elution profiles of frozen P-Rex2 DH/PH without CPA (magenta) and with 5% sucrose (peach) compared to a previously published freshly purified P-Rex2 DH/PH sample (Anderson et al. 2026; SASDB#). B-C. Guinier analyses and normalized residuals of frozen P-Rex2 DH/PH samples.
